# CRISPRi screens identify the lncRNA, *LOUP,* as a multifunctional locus regulating macrophage differentiation epigenetically and inflammatory signaling through a short, encoded peptide

**DOI:** 10.1101/2023.12.19.572453

**Authors:** Haley Halasz, Eric Malekos, Sergio Covarrubias, Samira Yitiz, Christy Montano, Lisa Sudek, Sol Katzman, S John Liu, Max A. Holbeck, Jonathan S Weissman, Susan Carpenter

**Affiliations:** Department of Molecular, Cell and Developmental Biology, University of California Santa Cruz, California, USA; Department of Biomolecular Engineering, University of California Santa Cruz, California, USA; Genomics Institute, Santa Cruz, CA 95064; Department of Radiation Oncology, University of California, San Francisco, San Francisco, CA 94158, USA Department of Neurological Surgery, University of California, San Francisco, San Francisco, CA 94158, USA; Department of Pediatrics, Division of Genetics and Genomics, Boston Children’s Hospital, Boston MA 02115; Department of Stem Cell and Regenerative Biology, Harvard University, Cambridge, MA, 02138 USA; Whitehead Institute for Biomedical Research, Massachusetts Institute of Technology, Cambridge, MA 02142, USA; Howard Hughes Medical Institute, Chevy Chase, MD 20815, USA; David H. Koch Institute for Integrative Cancer Research, Massachusetts Institute of Technology, Cambridge, MA 02142, USA; Department of Biology, Massachusetts Institute of Technology, Cambridge, MA 02142, USA

## Abstract

Long non-coding RNAs (lncRNAs) account for the largest portion of RNA from the transcriptome, yet most of their functions remain unknown. Here we performed two independent high-throughput CRISPRi screens to understand the role of lncRNAs in monocyte function and differentiation. The first was a reporter-based screen to identify lncRNAs that regulate TLR4-NFkB signaling in human monocytes and the second screen identified lncRNAs involved in monocyte to macrophage differentiation. We successfully identified numerous novel non-coding and protein-coding genes that can positively or negatively regulate inflammation and differentiation. To understand the functional roles of lncRNAs in both processes, we chose to further study the lncRNA *LOUP* (lncRNA originating from upstream regulatory element of *SPI1* [also known as PU.1]), as it emerged as a top hit in both screens. Not only does *LOUP* regulate its neighboring gene, the myeloid fate determining factor *SPI1*, thereby affecting monocyte to macrophage differentiation, but knockdown of *LOUP* leads to a broad upregulation of NFkB-targeted genes at baseline and upon TLR4-NFkB activation. *LOUP* also harbors three small open reading frames (sORFs) capable of being translated and are responsible for *LOUP*’s ability to negatively regulate TLR4/NFkB signaling. This work emphasizes the value of high-throughput screening to rapidly identify functional lncRNAs in the innate immune system.

## Introduction

According to the latest Gencode release (version 43), the human genome encodes 19,928 long noncoding RNAs (lncRNAs) making it the largest group of genes produced from the genome. Due to their cell type specificity this number continues to increase as more sequencing is performed (1). LncRNAs are transcripts over 200 nucleotides that are often spliced and polyadenylated and without annotated or predicted protein coding potential. Over the last decade a number of lncRNAs have been functionally characterized and shown to play diverse roles in a variety of biological processes from cell differentiation and cancer to immunity (2–4). Yet the functions of the vast majority of these transcripts remains unknown. Historically one of the largest challenges in studying lncRNAs has been the lack of reliable and specific approaches to target these transcripts, especially in a high-throughput manner. Because lncRNAs lack open reading frames, these genes are not susceptible to frameshift mutations induced by classic CRISPR/Cas9. Recently the adoption of the modified CRISPR/Cas9 technology -CRISPR inhibition (CRISPRi), has become a powerful tool for interfering with transcription of lncRNAs by inducing repressive chromatin marks at the transcription start site, making it an attractive approach to discover functional lncRNAs. Advanced computational developments in sgRNA library design coupled with targeted transcriptional repression induced by the components of CRISPRi, have made it possible to rapidly identify many functional lncRNA loci in a single pooled screening experiment (5).

A small number of high throughput screens have been performed to identify functional lncRNAs (5–8) but very few have been performed in immune cells (9, 10). To obtain new insights into innate immunity, more specifically monocyte and macrophage biology, we conducted two pooled CRISPRi screens in human monocytic cells (THP1s). In one screen, we sought to identify lncRNAs that regulate monocyte to macrophage differentiation, while the second assessed for lncRNAs that modulate NFkB signaling. Monocyte to macrophage differentiation is a highly coordinated process crucial to a well-regulated inflammatory response (11, 12). Acute activation of NFkB signaling in monocytes is imperative for proper recognition and resolution of pathogens. Therefore, it is important that we gain a more complete molecular understanding of both monocyte differentiation and NFkB regulation, as dysregulation of either of these processes can lead to a diseased state (13, 14). Acute activation of NFkB signaling in monocytes is imperative for proper recognition and resolution of pathogens, therefore it’s important that we gain a more complete molecular understanding of both monocyte differentiation and NFkB regulation, as dysregulation of either of these processes can lead to a diseased state (13, 14). In order to conduct the two screens, we developed a sgRNA library targeting over 2,000 THP1-expressed lncRNAs. Utilizing our newly developed NFkB (RelA/p65) reporter THP1 CRISPRi line we identified 35 regulators of NFKB. We utilized phorbol esters (PMA) to initiate differentiation and identified 38 lncRNAs regulating monocyte to macrophage differentiation. We were intrigued to find that one lncRNA, *LOUP*, was a top hit in both screens. Interestingly, *LOUP* neighbors the gene SPI1, a myeloid fate determining transcription factor required for development of monocytes and macrophages. Since many functional lncRNAs exhibit cis-regulatory effects, including *LOUP* (15), we sought to further investigate *LOUP*’s role as a cis regulator of *SPI1*. While Trinh et al., previously demonstrated that *LOUP* RNA mediates direct interactions with the *SPI1* promoter and the transcription factor RUNX1 in unstimulated monocytes, we found that *LOUP* and *SPI1* occupy a topologically associating domain (TAD) that is maintained in THP1s before and after differentiation, and found evidence of a prominent super enhancer (SE) in this TAD (16). The TAD and super enhancer are conserved in mouse bone marrow derived macrophages (BMDMs) and dendritic cells (BMDCs), highlighting the cross-species importance of cis-regulatory activity and chromatin structure at this locus in monocytes and their derivative cell types.

While we were able to verify that *LOUP* exhibits a cis-regulatory effect on *SPI1*, this did not account for its role as a negative regulator of NFkB. More recent discoveries in the field of lncRNA biology have uncovered lncRNAs that are bifunctional, where the RNA transcript can carry out *cis* or *trans* effects, but can also produce functional short, encoded peptides (SEPs) from short open reading frames (sORFs) (17). We analyzed ribosome sequencing (Ribo-Seq) and conservation data and found that *LOUP* is translated and harbors two sORFs well conserved in monkeys and apes. We conducted functional experiments that demonstrate that a *LOUP* SEP is responsible for *LOUP*’s negative regulation of inflammation. Together, this work defines an unappreciated role for this lncRNA in regulating critical immune processes, which was enabled by unbiased high throughput CRISPRi screening.

## Results

### CRISPRi screen identifies lncRNAs that regulate NFkB and macrophage differentiation

To begin to define lncRNAs that regulate NFkB signaling in human macrophages, THP1 cells containing five NFkB (p65) binding sites upstream a minimal CMV-driven EGFP, as well as deactivated Cas9-KRAB (dCas9-KRAB), were transduced with pooled lentivirus (MOI = 0.3) generated from our custom sgRNA library containing ∼25,000 individual sgRNAs (Table S1). The sgRNA library was designed using the hCRISPRi-v2.1 algorithm (5), with 10 sgRNAs targeting the transcription start sites of 2,342 lncRNAs annotated in the human genome assembly GRCh37 (hg19). LncRNA sgRNA target sites were determined based on expression in RNA-seq data from THP1s generated by our lab and previously published p65 ChIP-seq data and fantom transcription start site information (18) (Fig. 1A). The same design and cloning strategy were used as previously described (5).

**Figure 1.**
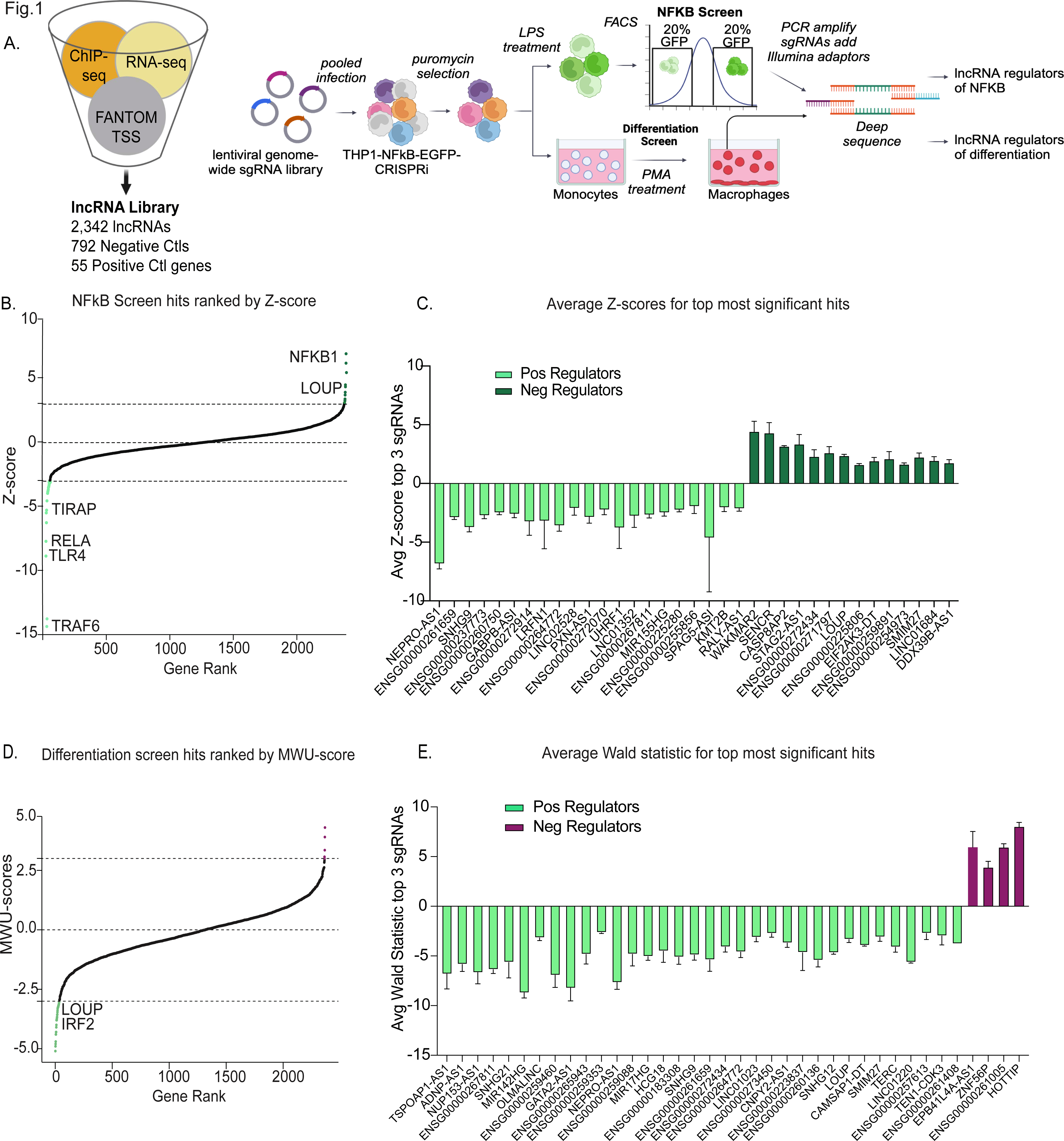
- CRISPRi screens identify positive and negative regulators of NFkB and macrophage differentiation. **A. Overview of sgRNA library design and overview of the screen.** sgRNAs were designed to target the transcription start sites of over 2000 Gencode hg19 annotated lncRNAs. Transcription start sites were predicted using data from FANTOM and ENCODE. THP1 lncRNA expression was estimated from THP1 RNAseq data. NFkB-EGFP-CRISPRi-THP1 cells were infected with pooled sgRNA libraries, selected, stimulated with LPS, and then sorted based on the top and bottom 20% of EGFP fluorescence. sgRNAs from the resulting GFP expressing populations were PCR amplified and sequenced. The same untreated THP1s containing the sgRNA library were treated with PMA and undifferentiated cells were collected 11 days post-treatment. sgRNAs from each timepoint were PCR amplified and sequenced. **B. NFkB screen analysis.** Analysis was performed on each of 3 screen replicates comparing sgRNA enrichment in the GFP low population or the GFP high population to the unsorted population with the MAUDE tool. Z-scores across replicates were combined by Stouffer’s method and ranked in order of Z-score. Z-score cutoffs of -3 and 3 are considered significant as highlighted in light green (positive regulators) and dark green (negative regulators). Genes labeled are significant known protein coding regulators of NFkB that acted as positive controls in the screen. **C. Significant NFkB hits.** The average z-scores of the top three best scoring sgRNAs for all significant hits with standard deviation. **D. PMA screen analysis.** DESeq2 was used to establish log2foldchange (L2FC) of sgRNAs between no treatment and PMA conditions. L2FC for sets of sgRNAs targeting each gene were compared to L2FC of all negative controls by Mann-Whitney U (MWU) test. MWU scores of -3 and 3 are considered significant. **E. Significant differentiation hits.** The average Wald test statistic of the top three best scoring sgRNAs for all significant hits with standard deviation

For the NFkB reporter screen, cells were stimulated with LPS for 24 h and then sorted by flow cytometry-assisted cell sorting (FACS). The top and bottom 20% of GFP+ gated cells were collected, and genomic DNA was isolated from each population (GFPhi and GFPlo). This gating strategy was determined based on earlier reporter-based sorting screens to sufficiently capture non-targeting controls (10, 18–20). Since lentiviral sgRNAs integrate into the genome, genomic DNA was isolated from each GFPhi and GFPlo and unsorted population. In each resulting population sgRNAs were PCR amplified and then sequenced (Fig 1A). MAUDE (21) analysis was performed comparing each GFPhi and GFPlo population to the unsorted (Fig. 1B). Genes with combined Z-scores (Stouffer’s method) of less than -3 were defined as significant positive regulators of NFkB while genes with combined Z-scores above 3 were defined as significant positive regulators (Table 1 and Table S2-3) Fig. 1B-C.

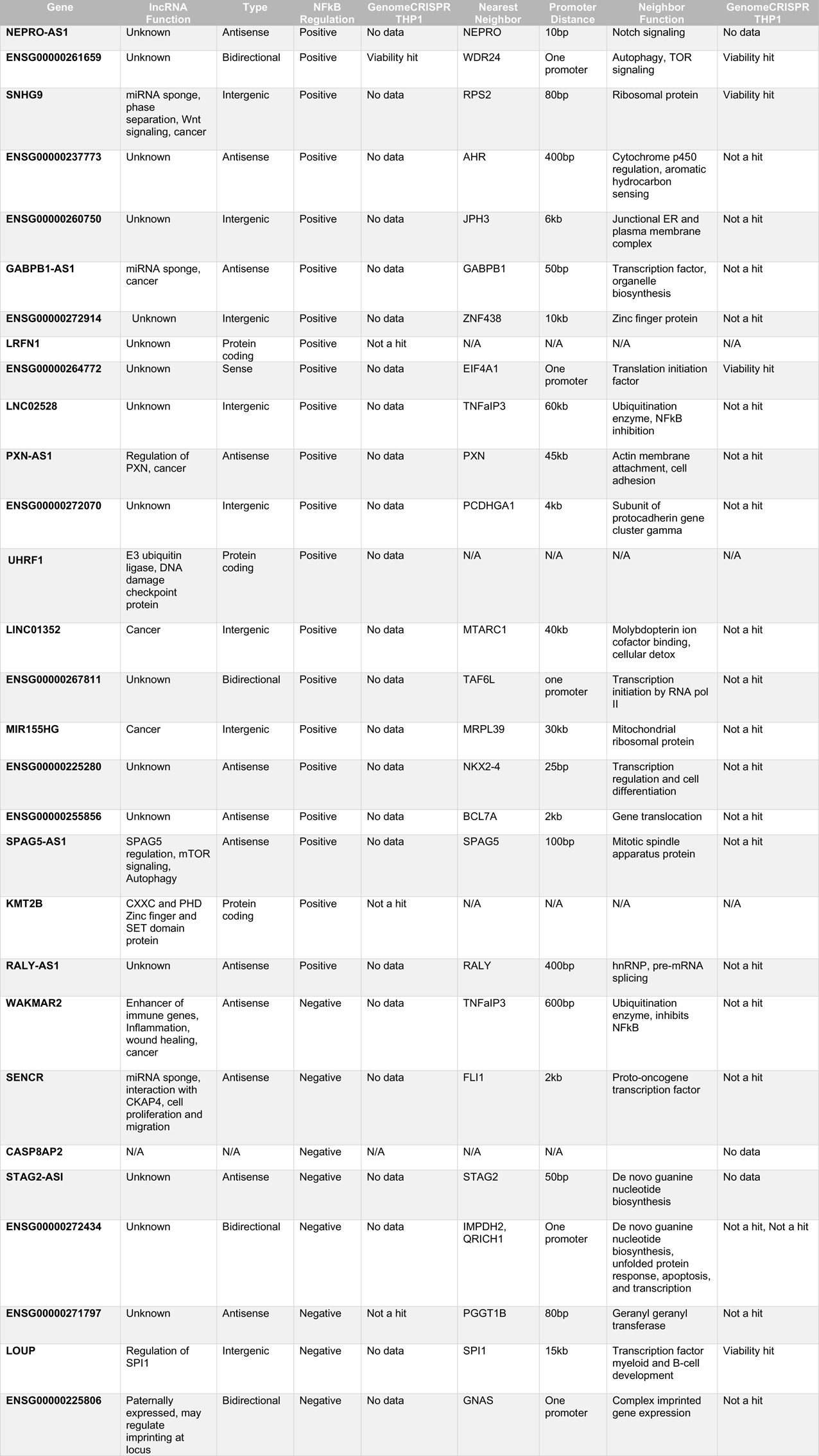
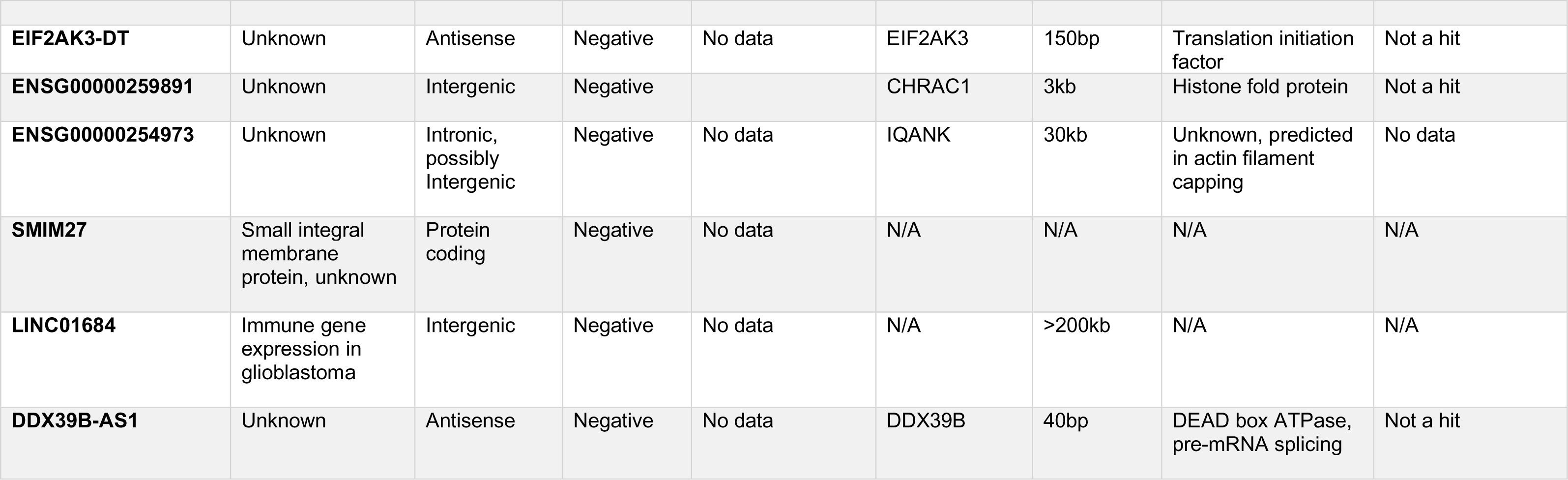

To screen for regulators for regulators of monocyte to macrophage differentiation. Following selection, cells were treated three times over the course of 11 days with 2nM PMA allowing for 50% of cells to differentiate (adhere to the plate), ensuring discovery of both positive and negative regulators of differentiation (Fig. 1A). 38 lncRNAs were identified with a Mann Whitney U-score greater than 3 or less than -3 (67 had a p-value cut off of < 0.01) (Table 2 and Table S4-6) as regulators of differentiation with over 75% representing negative regulators of the pathway (Fig. 1D-E).

| Gene | Function | Type | Differentiation Regulation | GenomeCRISPR THP1 | Nearest Neighbor | Promoter Distance | Neighbor Function | GenomeCRISPR THP1 |
| --- | --- | --- | --- | --- | --- | --- | --- | --- |
| TSPOAP1-AS1 | Unknown, breast cancer risk | Antisense | Positive | No data | SUPT4H1 | 14kb | mRNA processing, transcription elongation | Viability hit |
| ADNP-AS1 | Unknown | Bidirectional | Positive | No data | ADNP | One promoter | Cancer proliferation, Wnt signaling, neuronal differentiation | Not a hit |
| NUP153-AS1 | Unknown | Antisense | Positive | No data | NUP153 | 500bp | Component of nuclear pore complex | Viability hit |
| ENSG00000267811 | Unknown | Bidirectional | Positive | No data | TAF6L | One promoter | Component of PCAF complex | Not a hit |
| SNHG21 | Unknown | Intergenic | Positive | N/A | FSD2 | 40kb | Unknown | Not a hit |
| MIR142HG | N/A | miRNA, intronic | Positive | N/A | TSPOAP1-AS1 | One promoter | lncRNA | No data |
| OLMALINC | Unknown | Intergenic | Positive | No data | SCD | 25kb | Lipid biosynthesis | Viability hit |
| ENSG00000259460 | N/A | Antisense | Positive | No data | MEIS2 | 2kb | Transcription regulator | Not a hit |
| GATA2-AS1 | Cancer | Antisense | Positive | No data | GATA2 | 500bp | Endothelial cell transcription factor | Not a hit |
| ENSG00000265943 | Unknown | Antisense | Positive | No data | RBBP8 | 200bp | Nuclear protein binds RB and regulates cell proliferation | Viability hit |
| ENSG00000259353 | Unknown | Antisense | Positive | No data | SLTM | 13kb | Enables RNA binding activity | Not a hit |
| NEPRO-AS1 | Unknown | Bidirectional | Positive | No data | NEPRO | One promoter | Notch signaling | No data |
| ENSG00000259088 | Unknown | Antisense | Positive | Not a hit | PPP2R5C | 500bp | Negative control of cell growth and division | Not a hit |
| MIR17HG | Cancer, cell proliferation | miRNA | Positive | N/A | GPC5 | 50kb | Cell surface proteoglycan | Not a hit |
| HCG18 | Cancer, cell proliferation | Antisense | Positive | No data | TRIM39 | 500bp | Subunit of ORC complex, initiation of DNA replication | Not a hit |
| ENSG00000183308 | Unknown | Antisense | Positive | No data | ORC2 | 500bp | Subunit of ORC complex, initiation of DNA replication | Not a hit |
| SNHG9 | Cancer, cell proliferation | Bidirectional | Positive | No data | RPS2 | One promoter | Component of the ribosome | Viability hit |
| ENSG00000261659 | Unknown | Bidirectional | Positive | Viability hit | WDR24 | One promoter | Component of GATOR complex, TORC1 signaling pathway | Viability hit |
| ENSG00000272434 | Unknown | Bidirectional | Positive | No data | IMPDH2 | One promoter | Regulation of cell growth | Not a hit |
| ENSG00000264772 | Unknown | Sense overlapping | Positive | No data | EIF4A1 | One promoter | Enables double-stranded RNA binding activity | Viability hit |
| LINC01023 | Regulator of AKT in glioma, Hepatoblastoma tumorigenesis | Intergenic | Positive | No data | FER | 20kb | Tyrosine kinase, cell surface/ cytoskeleton signaling | Not a hit |
| ENSG00000273450 | Unknown | Intronic | Positive | No data | PLCE1 | N/A | Phospholipase that generates IP3 and DAG | Not a hit |
| CNPY2-AS1 | Unknown | Antisense | Positive | No data | CNPY2 | 10kb | FGF signaling regulator | No data |
| ENSG00000223837 | Unknown | Intronic | Positive | No data | BRD2 | One promoter | BET transcriptional regulator, Mitosis | Not a hit |
| ENSG00000260136 | Unknown | Bidirectional | Positive | Not a hit | COG7 | One promoter | Golgi morphology and localization | Not a hit |
| SNHG12 | miRNA sponge, cancer, angiogenesis | Intergenic | Positive | No data | TAF12 | 10kb | Transcriptional control of RNA pol II | Not a hit |
| LOUP | Regulation of SPI1 | Intergenic | Positive | No data | SPI1 | 15kb | Transcription factor myeloid and B-cell development | Viability hit |
| CAMSAP1-DT | Unknown | Bidirectional | Positive | No data | CAMSAP1 | One promoter | Microtubule polymerization | Not a hit |
| SMIM27 | Small integral membrane protein, unknown | Protein coding | Positive | No data | N/A | N/A | N/A | N/A |
| TERC | Telomerase RNA component | Intergenic | Positive | No data | N/A | N/A | N/A | N/A |
| LINC01220 | Carcinoma cell proliferation | Intergenic | Positive | No data | FOS | 15kb | Leucine zipper that dimerizes with JUN proteins to form AP-1 transcription factor | Not a hit |
| ENSG00000257613 | Unknown | Intronic | Positive | No data | PRANCR | One promoter | lncRNA regulator of epidermal homeostasis | No data |
| TEN1 | Telomere associated complex subunit | Protein coding, bidirectional | Positive | Not a hit | ACOX1 | One promoter | Fatty acid beta-oxidation | Not a hit |
| EPB41L4A-AS1 | Cancer, cell proliferation | Antisense | Negative | N/A | EPB41L4A | >200kb | Regulates interactions between the cytoskeleton and plasma membrane | Not a hit |
| ZNF506 | DNA binding transcription factor activity | Protein coding | Negative | Not a hit | N/A | N/A | N/A | N/A |
| ENSG00000261005 | Unknown | Intergenic | Negative | N/A | MLLT6 | 3kb | Histone binding activity, nucleosome binding activity | Not a hit |
| HOTTIP | Hepatocellular carcinoma, binds to WDR5 | Antisense | Negative | No data | HOXA13 | One promoter | Transcription factor | Not a hit |

Given that our sgRNA library included lncRNA genes regardless of genomic location, many of the targets are very near to or overlapping with neighboring genes. Heterochromatin induced by the dCas9-KRAB has been shown to reach as far as 1kb so we considered this caveat when choosing a top candidate lncRNA to further functionally validate (5, 22). 10 of the top 35 NFkB hits and 8 of the top 38 PMA hits are intergenic; defined as having their own promoters at least 1kb away from promoters of neighboring genes (Tables 1-2). 3 top NFkB hits, *LRF1*, *UHRF1*, and *KMT2B*, were previously annotated as non-coding transcripts on hg19 but have now been annotated as coding genes with unknown functions (Table 1). The other top hits are annotated either from a bidirectional promoter or antisense and overlapping to coding genes. In these cases, it is possible that the dCas9-KRAB has disrupted transcription of both bidirectional and overlapping transcripts. Interestingly, none of the coding genes targeted have been previously identified as regulators of NFkB or monocyte differentiation and represent novel coding regulators of the pathway and therefore are still of interest (Tables 1-2). Surprisingly, 7 lncRNAs and one protein coding gene (*ENSG00000267811*, *Nepro*-*AS1*, *SNHG9*, *ENSG00000261659*, *ENSG00000272434*, *ENSG00000264772*, *LOUP*, and *SMIM27*) were hits in both screens (Tables 1-2), suggesting genes playing multiple functions in monocyte/macrophage biology but only *LOUP* is intergenic and neighbors a gene known to regulate myeloid differentiation (SPI1).

### *LOUP* is conserved and regulates its neighbor SPI1

Given that *LOUP* (lncRNA originating from the upstream regulatory element of *SPI1*) arose as a top hit in both screens and is intergenic we wished to determine mechanistically how it can be involved with two different biological process in monocytes. Given the recent evidence that *LOUP* acts as an enhancer for its neighboring gene *SPI1* (15), and the evidence of a conserved SPI1 super enhancer (SE) spanning the locus in human and mouse (16), we examined H3K27Ac ChIP-Seq in THP1s (23). ATAC-seq showed open chromatin at *LOUP* and ChIP-Seq analysis found broad deposition of H3K27Ac marks indicating that the *LOUP* locus meets the more expansive SE criteria (24) (Fig. 2A). Given the evidence of an important SE, we hypothesized that local chromatin structure would have important regulatory functions and called TADs from Hi-C data (23). TAD calling and CTCF ChIP-seq indicate that *LOUP* and *SPI1* occupy the same domain (Fig. 2A). GTEx gene expression shows that *LOUP* and *SPI1* are expressed almost exclusively in blood cells in contrast to neighboring genes outside the TAD, demonstrating that they are members of a distinct domain (Supp Fig 1).

**Figure 2.**
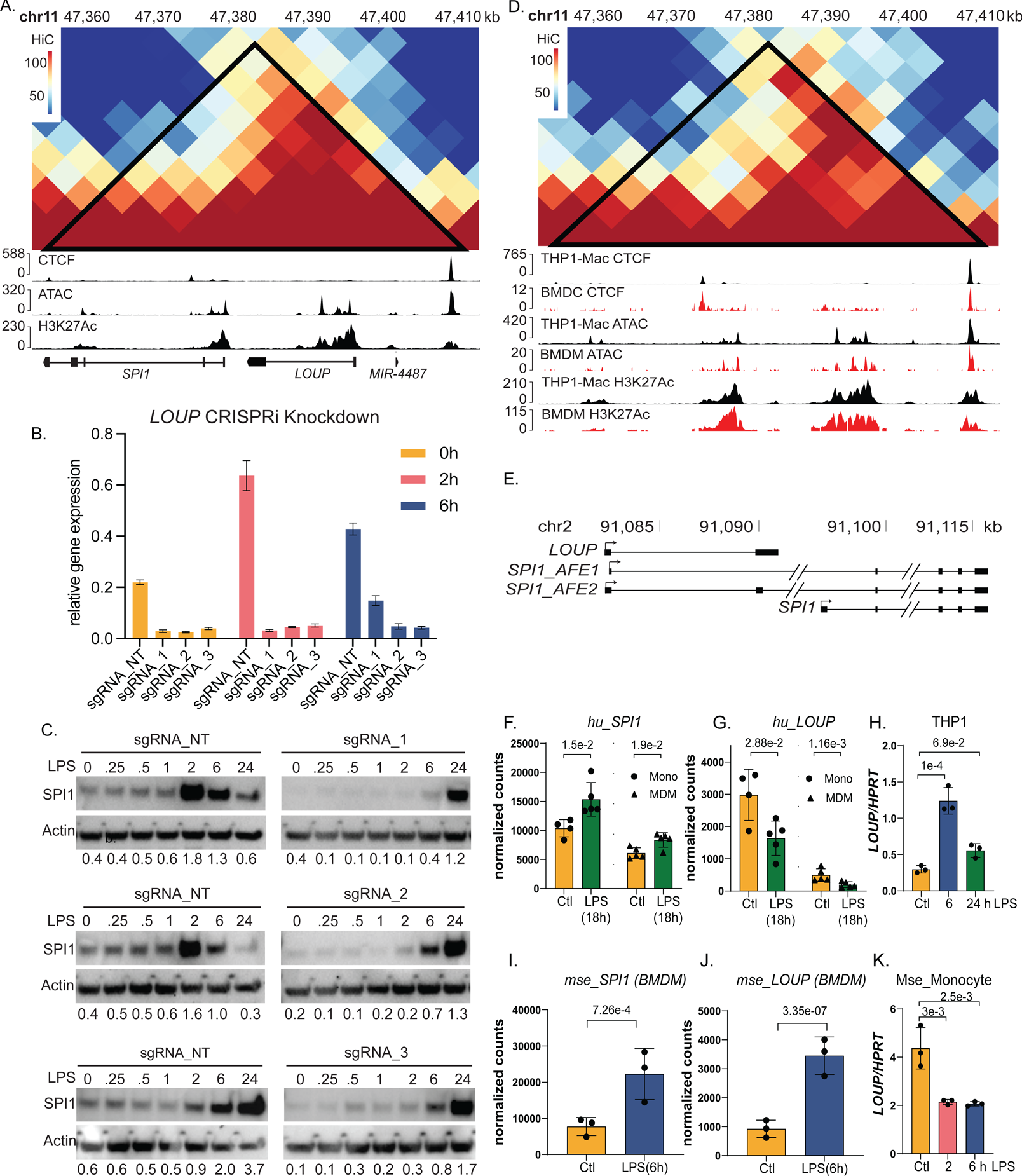
- *LOUP* regulates SPI1 in monocytes. **A. Predicted TAD for LOUP and SPI1 in THP1s**. Browser tracks display regions of CTCF binding, ATAC peaks, and regions of high H3K27 acetylation. Numbers on the Y-axis represent raw read counts. **B. CRISPRi Knockdown of *LOUP* in THP1s**. Three additional sgRNAs (sgRNAs 1, 2, 3) were designed to target top lncRNA candidate *LOUP*. qPCR measurement of *LOUP* across three replicate experiments shows knockdown of *LOUP* by all three sgRNAs (p-values<0.05) vs a non-targeting control sgRNA (NT) before LPS stimulation (0 hours) and after 2 and 6 hours of LPS stimulation. Values are normalized to HPRT and error bars represent standard deviation. **C. Western blot analysis of SPI1.** Non-targeting control (NT) vs. 3 *LOUP* knockdowns (sgRNAs 1, 2, 3) over a time course (hours) of LPS treatment. Samples were collected at baseline (0), 15min (.25), 30min (.5), 1 hour (1), 2 hours (2), 6 hours (6) and 24 hours (24). Quantitative values indicate densitometry ratios of SPI1:Actin. **D. Conserved TAD and epigenetic marks following differentiation in human and mouse.** PMA differentiated THP1-Macrophages (THP1-Macs) maintain chromatin structure and epigenetic marks at the *LOUP*/*SPI1* locus. Patterns of epigenetic marks and CTCF binding are also conserved in mouse BMDMs and BMDCs. **E. Loup and Spi1 isoforms in mouse.** Gene models of transcripts at Spi1 locus determined from long read Nanopore on BMDMs and called with FLAIR. **F-K. Effect of LPS Stimulus on *LOUP* and *SPI1* Expression in Human and Mouse Cells. F&G. *SPI1*(F) and *LOUP*(G) expression in Human Monocytes and Monocyte Derived Macrophages.** Differential gene expression was calculated from RNA-Seq of untreated or LPS treated monocytes and macrophages. Significance represented by adjusted p-value of DESeq2 implementation of the Wald-test. **H. qPCR of *LOUP* in THP1** qPCR measurement of *LOUP* across three replicates of Human THP1 cells untreated or treated with LPS for 6 or 24 hours. *LOUP* expression increases at 6 hours of LPS treatment and comes back down to a non-significant change compared to control by 24 hours. Values are normalized to HPRT. Error bars show standard deviation and significance testing was performed with one-way anova with Dunnett’s multiple comparisons in GraphPad Prism. **I&J. *Spi1*(I) and *Loup*(J) expression in Mouse Bone Marrow Derived Macrophages**. Differential gene expression was calculated from RNA-Seq of untreated or LPS treated BMDMs. Significance represented by adjusted p-value of DESeq2 implementation of the Wald-test. **K. qPCR of *LOUP* in Mouse Monocytes** qPCR measurement of *LOUP* across three replicates of primary mouse monocytes untreated or treated with LPS for 2 or 6 h. *LOUP* expression decreases in both LPS treated conditions relative to untreated control. Values are normalized to HPRT. Error bars show standard deviation and significance testing was performed with one-way ANOVA with Dunnett’s multiple comparisons in GraphPad Prism.

Given the recent evidence that *LOUP* may act as an enhancer for its neighboring gene *SPI1* (15), and that this is true for other lncRNAs neighboring critical protein regulators such as the lncRNA *lincRNA-Cox2* and PTGS2 (25), and *lncRNA-p21* and its protein neighbor p21 (26), we wanted to measure levels of SPI1 protein following *LOUP* knockdown. To this end we generated three independent sgRNA lines targeting *LOUP* and confirmed knockdown by qPCR compared to a scrambled control line (Fig. 2B). Next, we stimulated cells with 200ng/ml LPS for the outlined time course and measured SP1 levels by western blot (Fig. 2C). In THP1 cells SPI1 is present at baseline and is strongly induced between 2 and 6 h following LPS stimulation in non-targeting control cells (Fig. 2B). Perhaps most interesting is the comparison of SPI1 levels between each of the control replicates and three different *LOUP* knockdowns at baseline where SPI1 is downregulated when *LOUP* is knocked down. In the *LOUP* knockdown lines SPI1’s induction peaks after 24 h of LPS to the levels observed in the control lines after 2 to 6 h, indicating that *LOUP* acts as an enhancer for SPI1 but *LOUP* alone is not essential for SPI1 expression (Fig. 2C).

Since *LOUP* was a hit in the PMA differentiation screen, we performed epigenetic analysis on data from PMA differentiated THP1s. We found the Hi-C, ChIP-seq and ATAC-seq patterns to be consistent after PMA treatment, indicating that local epigenetic changes are not primarily responsible for differences in *SPI1* or *LOUP* expression following differentiation. Epigenetic marks are also consistent in mouse BMDMs and BMDCs compared to human macrophages indicating that the epigenetic landscape is strongly conserved in differentiated monocyte-derived cells across species (Fig. 2D). Despite this remarkable level of conservation, we find evidence that the *LOUP* transcript itself has diverged between human and mouse. Whereas human *LOUP* is a distinct gene, in mice we find that the equivalent locus is transcribed as either a two exon lncRNA (*Loup*) or as an extended 5’-UTR of Spi1 (Fig. 2E). There is no evidence of an extended 5’UTR of Spi1 in our human long read data (27). Moreover, the *LOUP*/*Loup* sequences are only 37% identical and have dissimilar predicted secondary structures (Supp Fig.2).

We next examined the expression levels of *LOUP* and *SPI1* across human and mouse monocytes and macrophages at baseline and following LPS stimulation. Interestingly *SPI1* and *LOUP* are dominantly expressed in primary monocytes (Mono, Fig. 2F and G left panels) compared to differentiated macrophages. *SPI1* is consistently induced following LPS (Fig. 2F and I), while *LOUP* appears to be reduced in expression in primary human and mouse monocytes (Fig. 2G and K) but induced in human monocytic cell line THP1 and mouse macrophages (Fig. 2H and J). These differences in expression patterns of *LOUP*/*Loup* might be explained by the complexity of the locus and the possible isoforms being produced as outlined in Fig. 2D and Supp Fig. 3

**Figure 3.**
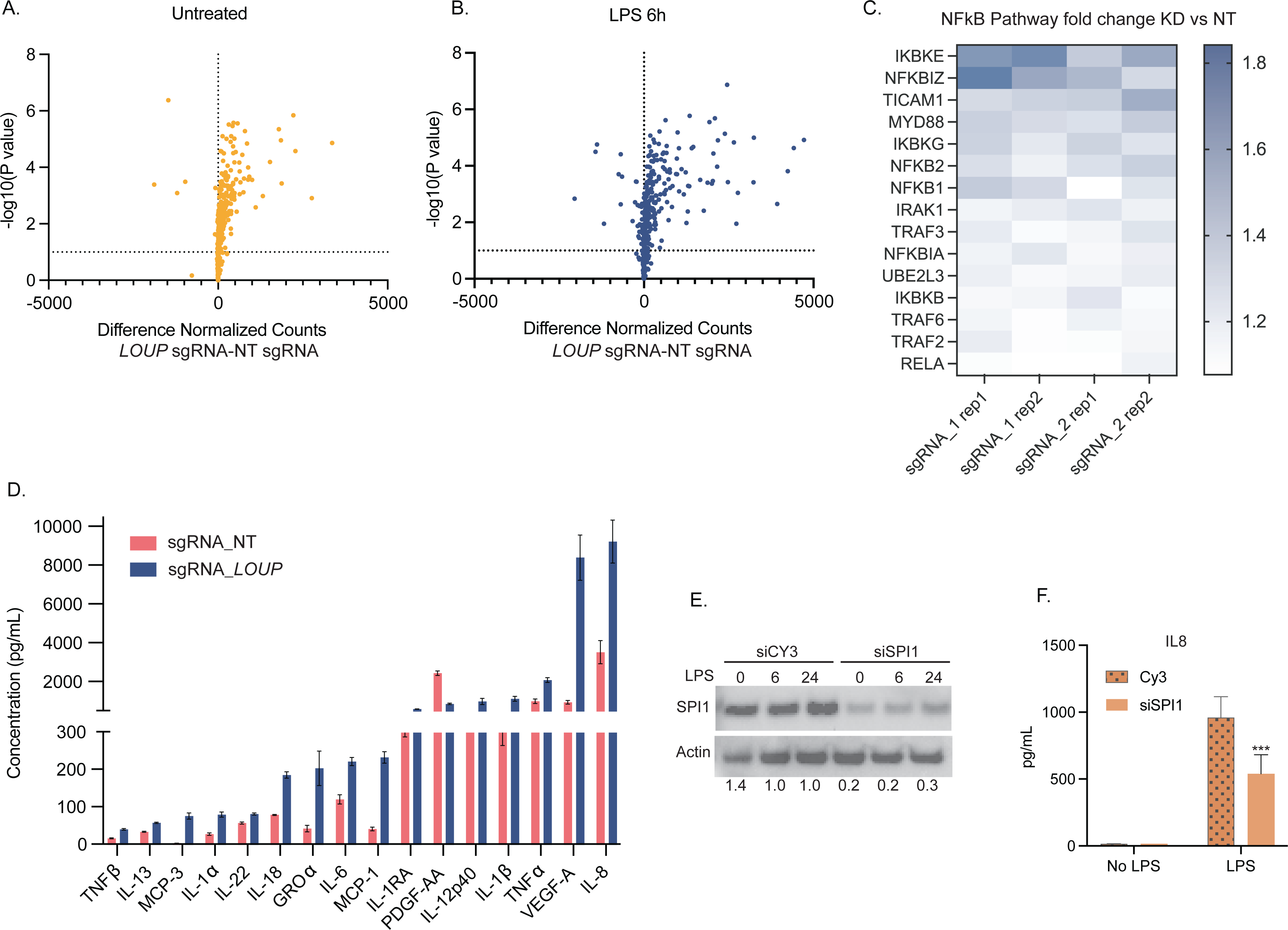
- Knockdown of *LOUP* upregulates NFkB targeted genes. **A-C. Multiplexed analysis of immune related transcripts upon *LOUP* knockdown. A-B**. Transcripts of 580 immune genes were measured in RNA from a non-targeting control and 2 of the sgRNA *LOUP* Knockdowns at baseline and after 6 hours of LPS stimulation. Each sample was measured in duplicate. All transcript counts were normalized to 6 housekeeping genes then both knockdowns and duplicate measurements were averaged. All genes were plotted regardless of p-value. **C**.. Heat map representing fold change of knockdowns vs non-targeting control at baseline for NFkB pathway genes significantly up-regulated in transcript analysis (A) (p-values<0.05). **D. Multiplexed ELISA analysis of cytokines upon *LOUP* knockdown.** ELISA was performed on the supernatant from three non-targeting controls and all three *LOUP* knockdowns after 24 h LPS treatment. All bars represent an average of all three non-targeting (NT) or an average of all three knockdowns with standard deviation. All differences observed between NT and knockdowns for each cytokine are significant (p-values<0.05). **E. Knockdown of SPI1.** Western blot analysis of SPI11 in WT THP1 cells transfected with a *SPI1* targeting siRNA (siSPI1) compared to a control non-targeting siRNA (siCy3). Samples were collected at baseline (0) and after 6 and 24 h of LPS treatment. Quantitative values indicate densitometry ratios of SPI1:Actin. Blot is representative of 3 separate experiments. **F. Effect of SPI1 knockdown on IL8.** IL8 ELISA was performed on supernatant from control and *LOUP* siRNA THP1s treated with LPS for 24 h. Bars represent an average of two separate experiments each measured in triplicate with standard deviation (p-value<0.0005).

### *LOUP* acts to negatively regulate NFkB target genes at the RNA and protein level

Based on the results of the screen, *LOUP* can negatively regulate NFkB activation (Fig 1.B and C). We aimed to look broadly at inflammatory gene expression upon *LOUP* knockdown (LOUP-KD). RNA was collected from two of the *LOUP*-KD lines (sgRNA_1 and sgRNA_2) both before and after 6 h of LPS stimulation (Fig. 3A-B). Using Nanostring technology (Immune gene panel) to directly quantify RNA transcripts of over 500 immune genes, it was evident that knockdown of *LOUP* broadly upregulates transcription of inflammatory genes both before and after LPS stimulation, including transcripts that comprise the TLR4/NFkB signaling pathway (Fig. 3C). To determine if knockdown of *LOUP* affected protein levels of inflammatory genes, we collected supernatant 24 h post LPS treatment from all three *LOUP* knockdown THP1 cell lines and performed cytokine arrays testing 45 proteins. 16 of the 45 inflammatory cytokines were significantly increased compared to controls, with IL8 being the most upregulated (Fig 3.D).

To determine if a reduction in *SPI1* could be responsible for the negative regulation of NFkB or inflammatory signaling, THP1s were transfected with siRNA targeting *SPI1* or a non-targeting siRNA control (siCY3) and knockdown was confirmed by western blot (Fig. 3E). IL8 was measured in the *SPI1* knockdown vs control by ELISA. In contrast to *LOUP* knockdown, *SPI1* knockdown resulted in significantly decreased levels of IL8 (Fig. 3F). This is consistent with the role of *SPI1* as a positive regulator of inflammation (28–30) and suggests that loss of SPI1 upon *LOUP* knockdown is not responsible for the increase in inflammatory gene expression.

### A *LOUP* sORF encoded peptide (SEP) functions as a negative regulator of NFkB

To determine *LOUP*s mechanism of action in regulating inflammatory genes, we first determined its localization in macrophages following fractionation and qPCR. *LOUP* RNA was detected in both the nucleus and the cytoplasm (Fig. 4A). Nuclear localization is consistent with its roles in regulating transcription of *SPI1* and we reasoned that its dominant expression in the cytoplasm could explain the effects seen more broadly on gene targets of NFkB through the production of SEPs. It’s been found that some cytoplasmic lncRNAs harbor short open reading frames capable of producing peptides less than 100 amino acids in length, and that these peptides can carry out important functions (31). To investigate the coding potential of *LOUP* we first examined existing Ribo-seq datasets from THP1s and primary macrophages (Fig. 4B). Based on ribosome footprints we established that *LOUP* harbors three sORFs with potential for translation.

**Figure 4.**
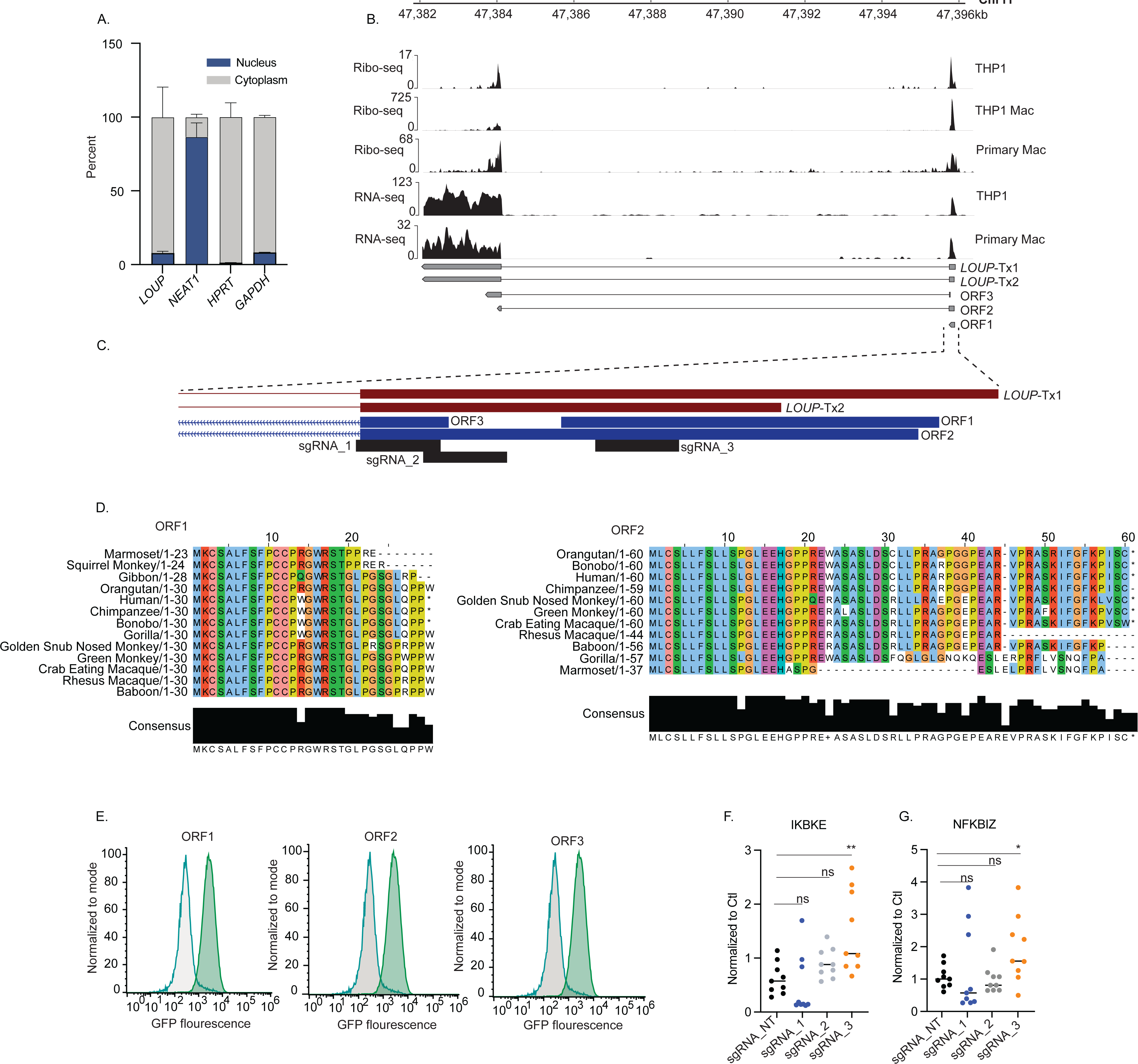
- *LOUP* encodes a SEP that regulates NFkB target genes. A. **Cellular localization of *LOUP* in THP1s.** Nuclear/cytoplasmic fractionation of WT THP1 cells after 6 hours of LPS stimulation. Data is from a single experiment with propagated error calculated for triplicate measurements. **B. *LOUP* sORF translation.** Annotations of two *LOUP* isoforms and of three short open reading frames (sORFs) predicted for LOUP gleaned from Ribo-seq tracks generated from THP1s, differentiated THP1s and primary macrophages. Also included are tracks for short read RNA-seq from THP1s and long read RNA-seq (R2C2) from primary macrophages. B. **Cas9 targeting of sORFs.** *LOUP* isoforms showing the use of two transcription start sites and the translation initiation positions of sORFs, followed by depiction of sgRNAs targeting sORFs with CRISPR-Cas9. sgRNAs 1-3 correspond to panel F. **D. Conservation of sORFs.** Human sORF sequences were aligned to genome assemblies of other primates with Blat, then sequences were translated. **E. Expression of *LOUP* ORF-GFP.** Flow cytometry measuring GFP expression in LOUP knockdown THP1s transfected with sORF-GFP fusion constructs. **F. sORF regulation of inflammatory genes.** Two of the top most upregulated genes in the LOUP CRISPRi knockdowns were measured in LOUP Cas9 sORF targeted cells. Expression of IKBKE and NFKBIZ were measured at baseline by qPCR in each of 3 Cas9 cell lines (sgRNA_1, sgRNA_2, sgRNA_3) along with a non-targeting control (sgRNA_NT) (ns = not significant, * = p-value<0.05, ** = p-value<0.001).

Of the two *LOUP* transcript models established from our published isoform-level transcriptome atlas of macrophages (27), only the longer transcript, LOUP-Tx1, contains all three ORFs, while the shorter transcript, LOUP-Tx2, is limited to ORF3 (Fig. 4C). We note that despite ORF1 initiating at the 5’-most ATG, the putative 5’UTR in this case is only 14 nucleotides and recent evidence suggests that such short UTRs can favor translation initiation from downstream start codons (32). Taken together, the evidence of translation from all three ORFs is consistent with different TSS usage and low initiation stringency from a short 5’UTR.

We next looked at conservation of the sORFs and found that ORF1 and ORF2 are highly conserved in primates (Fig. 4D) while ORF3 is not. There is no evidence of conservation of the ORFs in mouse. To test the translational potential of these three sORFs, each was cloned in frame with GFP to create a translational fusion and introduced into THP1-NFkB-CRISPRi cells. Measurable GFP expression was driven by all three sORFs (Fig 4E). Next, sgRNAs were designed to target the sORF regions in THP1s with active CRISPR-Cas9 (Fig. 4C). To determine if disruption of any sORFs resulted in an increase of inflammatory cytokines at baseline in accordance with the CRISPRi results, we measured transcript levels of *IKBKE* and *NFKBIZ* (Fig. 4F-G). These two transcripts were most significantly upregulated at baseline in the *LOUP* CRISPRi knockouts (Fig. 3C). Interestingly both genes were significantly upregulated by the sgRNA targeting ORF1 and ORF2, but not the sgRNA uniquely targeting ORF2 (Fig. 4D). Together this data confirms that *LOUP* is a bimodal locus capable of regulating its neighboring gene through an enhancer mechanism as well as encoding an SEP that functions to negatively regulate NFkB genes in monocytes.

## Discussion

Here we’ve described genome wide pooled CRISPRi screening to identify novel lncRNA genes that regulate NFkB signaling and monocyte to macrophage differentiation. NFkB signaling pathways have been extensively studied and protein coding genes that regulate the signaling cascades have been considered mostly resolved (14, 33–35). Not only have we successfully identified novel lncRNA regulators of the TLR4/NFkB pathway, but we have also unveiled novel protein regulators. LncRNAs that share a promoter or overlap with protein coding regions pose a great challenge for functional characterization. By targeting these genes with CRISPRi we may have simultaneously disrupted the function of the overlapping coding gene, regardless of how specifically targeted the dCas9-KRAB was to the lncRNA TSS. Despite this caveat, we have identified novel coding genes that function in this context. Further studies using classic CRISPR Cas9 to disrupt coding potential of these genes and measure effects on NFkB signaling are warranted to confirm that it is indeed the protein mediating the phenotype (Tables 1-2). Although phenotypes associated with lncRNAs are often subtler than those of proteins, we’ve used stringent analysis thresholds to bolster confidence in our list of functional lncRNA candidates. One candidate in particular, *LOUP*, that has been previously identified as an enhancer lncRNA in myeloid cells (15), served as an excellent target for further functional validation. Not only were we able to confirm its ability to function as an enhancer in a monocyte cell line and impact the differentiation process, but we’ve also begun to unravel its novel mechanistic role as a negative regulator of TLR4/NFkB signaling.

It is known that activation of TLR4 signaling can prime defined enhancer regions in both human and mouse macrophages (17, 36). In addition to the specific histone modifications that define enhancer regions, the actual transcription of enhancer RNAs from these regions (eRNAs) seems to be an important mechanism for enhancer function during TLR4/NFkB-driven gene expression (36). Although the question remains whether production of eRNAs are simply a consequence of transcriptional regulation by these loci or if these eRNA transcripts themselves are functional. It has been established that many lncRNA genes can also exert transcriptional regulation on nearby genes without having the definitive histone marks that delineate an enhancer (37, 38). Cis upstream regulatory elements of *SPI1* have also been previously identified (39–42), but more recently it has been shown that the functional and myeloid-specific lncRNA *LOUP*, is transcribed from an upstream element of SPI1 (15). The authors demonstrated that the lncRNA transcript itself mediates the interaction between the *SPI1* promoter and the transcription factor RUNX1, but that the *LOUP* gene does exhibit histone modifications consistent with an enhancer (high H3K4me1, low H3K4me3). While we first identified *LOUP* for its ability to regulate the activity of TLR4/NFkB signaling and monocyte differentiation in our screens, we were able to confirm the previous finding that it acts as an enhancer of *SPI1*. *SPI1* is a myeloid lineage-determining transcription factor that is also active in the inflammatory response of monocytes and macrophages (28–30). In addition to the fact that *LOUP* and *SPI1* occupy the same TAD, we’ve shown that *SPI1* is induced in monocytes (Fig. 2C) upon treatment with LPS and that absence of *LOUP* decreases *SPI1* expression, especially at baseline. Interestingly, expression of *SPI1* in these monocytes seems to be able to overcome the absence of *LOUP* by 24 h, perhaps indicating the essential role of the protein in these cells, and that there are multiple routes that can lead to its regulation (39, 43, 44).

The essential role of SPI1 in monocytes and monocyte-derived cells is underscored by the striking conservation of H3K27Ac marks, open chromatin, and CTCF binding at the locus in both human and mouse (Fig. 2A,D). Despite this, *LOUP* itself exhibits the low cross-species conservation characteristic of many lncRNA genes. Whereas our mouse *Loup* long read data demonstrates that this locus can be transcribed as an independent lncRNA or as an alternate first exon for *Spi1*, we find no evidence for the latter in human *LOUP* (Fig 2E,4B). In the other direction, *LOUP* in primates has developed, or maintained, conserved, and translated sORFs that are missing in mouse (Fig. 4D). Taken together we report a remarkable combination of highly conserved regulatory elements and divergent transcript features at the loci in mouse and human.

Interestingly, we found that hundreds of inflammatory genes are upregulated upon suppression of the *LOUP* locus both at baseline and upon LPS stimulation. Of the secreted cytokines measured in the *LOUP* knockdowns (Fig 2.D), IL-8 was one of the most upregulated, but was significantly decreased in response to knocking down *SPI1*, indicating that *LOUP’s* effect on IL8 is independent of its role as an enhancer of *SPI1*. This was not surprising given SPI1’s known role in positively regulating expression of inflammatory genes. Given that *LOUP* was a top hit in our NFkB-reporter based screen, along with the fact that *LOUP* is present in the cytoplasm and harbors three small ORFs, led us to investigate the possibility that *LOUP* encodes a functional peptide capable of negatively regulating the NFkB-driven inflammatory response in these cells. Based on the association of *LOUP*’s sORFs with ribosomes together with the fact that targeting the sORF region using active CRISPR resulted in upregulation of inflammatory genes, we believe that *LOUP* produces a functional peptide. The minimum cutoff employed for annotated protein coding genes is typically 100 amino acids. It’s now appreciated that many annotated lncRNAs harbor sORFs that would result in peptides well below this cut-off (31). CRISPR-Cas9 disruption of ORFs 1 and 2, which are predicted to be 30 and 59 amino acids respectively, resulted in an increase in inflammatory gene expression, while disruption of ORF3, predicted to be 123 amino acids, did not have an effect (Fig 4F). While questions remain about the functionality of these peptides under 100 amino acids, there is mounting evidence suggesting that they have evolved to carry out important immunological functions (17, 45–47). In addition, there is evidence supporting the potential for both the lncRNA transcript itself along with lncRNA derived SEPs to have distinct functional roles. For instance, the lncRNA *Aw112010,* has been shown to downregulate the transcription of *IL10* through interactions with the histone demethylase KMD5, thereby controlling T-cell differentiation (48). *Aw112010* also produces an LPS-inducible SEP (84 amino acids) that regulates the mucosal inflammatory response in mice without effects on IL-10 (46). Our work here illustrates that a similar bimodal mechanism might be the case for the *LOUP* gene, in which the lncRNA regulates transcription of its neighboring gene in *cis* while the SEP modulates the NFkB-driven inflammatory response in *trans*. While we show *LOUP* produces a functional peptide, more work needs to be done to understand the mechanisms by which this peptide directly or indirectly impacts the expression of innate immune genes.

Here we showcase the value of pooled CRISPRi screening as an efficient method to identify functional lncRNAs in the context of innate immunity. Meticulous control of macrophage signaling is crucial for a proper immune response and the screens performed here have demonstrated that lncRNAs play an important role in maintaining these signaling pathways. Understanding lncRNA function in this context has led to novel insights into inflammatory gene regulation and even the bimodal functional capabilities of lncRNAs to regulate gene expression in *cis* and *trans*. Here we described a novel role for a myeloid-specific lncRNA as a potent regulator of inflammatory gene expression. This work will serve as a valuable resource of both lncRNAs and coding genes previously undiscovered in this pathway, as well as an important foundation for further mechanistic understanding of functional SEPs.

## Methods

### Cell lines

Wildtype (WT) THP1 cells were obtained from ATCC. All THP1 cell lines were cultured in RPMI 1640 supplemented with 10% low-endotoxin fetal bovine serum (ThermoFisher), 1X penicillin/streptomycin, and incubated at 37°C in 5% CO2.

#### Lentivirus production

All constructs were cotransfected into HEK293T cells with lentiviral packaging vectors psPAX (Addgene cat#12260) and pMD2.g (Addgene cat#12259) using Lipofectamine 3000 (ThermoFisher cat# L3000001) according to the manufacturer’s protocol. Viral supernatant was harvested 72 hours post transfection.

#### THP1-NFkB-EGFP-dCasKRAB

We constructed a GFP-based NF-κB reporter system by adding 5x NF-κB-binding motifs (GGGAATTTCC) upstream of the minimal CMV promoter-driven EGFP. THP1s were lentivirally infected and clonally selected for optimal reporter activity. Reporter cells were then lentivirally infected with the dCas9 construct that was constructed using Lenti-dCas9-KRAB-blast, addgene#89567. Cells were clonally selected for knockdown efficiency greater than 90%.

#### THP1-NFkB-EGFP-dCasKRAB-sgRNA (LOUP knockdown)

NFkB-EGFP-CRISPRi-THP1 cells were lentivirally infected with sgRNAs. sgRNA constructs were made from a pSico lentiviral backbone driven by an EF1a promoter expressing T2A flanked genes: puromycin resistance and mCherry. sgRNAs were expressed from a mouse U6 promoter. 20-Nucleotide forward/reverse gRNA oligonucleotides were annealed and cloned via the AarI site.

#### THP1-NFkB-EGFP-dCasKRAB-LOUP-/-sORF+

sORF-GFP fragments were synthesized by Twist Biosciences and cloned into a pSico lentiviral backbone. Constructs were then packaged into lentiviral particles as described above. Unstimulated GFP+ cells were sorted by FACS on a BD FACSAria II two times to achieve a 100% GFP positive population assuming that GFP expression in unstimulated cells was not activation of the reporter. Cells were consistently cultured under blasticidin and puromycin to maintain active dCas9 and sgRNA expression.

#### THP1-NFkB-EGFP-Cas9

The NFkB reporter was introduced as described above. The Cas9 construct was constructed from a pSico lentiviral backbone with an EF1a promoter expressing T2A flanked genes: blastocidin-resistant (blast), blue fluorescent protein, and humanized *Streptococcus pyogenes* Cas9.Cells were clonally selected for knockdown efficiency greater than 90%.

#### Primary Mouse Monocytes

Primary Mouse Monocytes were isolated by performing bone marrow extraction from 6 C57BL/6 mice per replicate followed by monocyte isolation using Monocyte Isolation Kit (Mouse) (Miltenyi, 130-100-629). Cells were cultured in DMEM media supplemented with 10% Heat inactiated Fetal Bovine Serum, 1X Pen Strep, 1X Cipro. Cell lines cultured at 37°C with 5% CO2.

### Screening protocol

#### sgRNA library design and cloning

10 sgRNAs were designed for each TSS of hg19 annotated lncRNAs expressed in THP1s at baseline and upon stimulation. The sgRNA library also included 700 non-targeting control sgRNAs, and sgRNAs targeting 50 protein coding genes as positive controls. The sgRNA library was designed and cloned as previously described in (5).

#### CRISPRi NFkB FACS Screen

Library infected and selected THP1-NFkB-EGFP-CRISPRi-sgRNA cells were expanded to 2000X coverage. Cells were stimulated with LPS (200 ng/mL) for 24 h to induce expression of NFkB-EGFP. Flow cytometry and PCR amplification of genomic sgRNA sequences were conducted as previously described in detail in (20). Screen was performed three times in THP1 lines (replicates A, B, C).

#### NFkB Screen Analysis

SgRNA guide adapters were removed with cutadapt (49) and counts were obtained with the MAGeCK count function from MAGeCK (50). Z-scores for each gene were found by analyzing each replicate independently with MAUDE (21). For each gene in the experiment, aggregate z-scores were generated using Stouffer’s method and a combined false-discovery rate was calculated.

#### CRISPRi PMA Screen

THP1-NFkB-EGFP-CRISPRi-sgRNA were infected with the sgRNA library as previously described for the NFkB screen. Cells line were infected and ∼7000X coverage was achieved. Triplicates were either left untreated or treated with 2nM PMA on days 0, 8, and 9. Undifferentiated cells were collected on day 11 and sgRNAs were PCR amplified as previously described for the NFkB screen.

#### PMA Screen Analysis

SgRNAs were counted and passed to DESeq2 for analysis. Default normalization was performed, and log2foldchange (L2FC) was calculated for each sgRNA between the PMA and No Treatment conditions. L2FC for the set of sgRNAs targeting each gene were compared to L2FC of all negative controls by Mann-Whitney U (MWU) test. PMA replicate B was excluded from the analysis as it fell below 500X sgRNA coverage over the course of the experiment.

#### Sequencing Data

Human RNA-Seq data are from (51) and available at GSE147310. We previously published the mouse BMDM RNA-Seq Data (52) and it is available at GSE141754. Data pertaining to ATACSeq, ChIPSeq and HiC analyses in THP1s were originally reported in (23) and are available at GEO: GSE96800 and SRA: PRJNA385337. RiboSeq data is from (53–55), with data available at GSE208041, GSE66810, GSE39561 respectively. BMDC CTCF ChIP-Seq is deposited at GSE36099 (56). BMDM ATAC-Seq and H3K27ac ChIP-Seq was used from the C57 strain deposited at GSE109965 (57).

#### ATAC and ChIP-Seq

Adapters were trimmed with Ngmerge (58) and mapped to GRCh38 primary assembly for human, or GRCm39 for mouse, with Bowtie2 (--very-sensitive --maxins 1000) (59). Replicates were merged and alignments were converted to BigWig tracks with the bamCoverage (--binsize 1) module from deepTools (60).

#### Liftover of mouse tracks

Mouse alignments were lifted to GRCh38 with the “liftover” utility from the UCSC Genome Browser’s kent tools using hg38.mm39.all.chain.gz. Alignments were visualized using pyGenomeTracks (61).

#### HiC

Paired-end reads from untreated or PMA treated THP1s were deduplicated with BBMap clumpify (dedupe=t ziplevel=3 reorder=t compresstemp=f deleteinput=t). Fastqs were converted to Pairs format with Chromap (-e 4 -q 1 --split-alignment --pairs) (62) and replicates were merged with Pairtools (63). Cool format files were generated with Cooler (cooler cload pairs) at 5kb resolution, then normalized with hicNormalize (-n smallest -sz 1.0) and Knight-Ruiz corrected with hicCorrectMatrix (--correctionMethod KR) (Abdennur and Mirny, 2020). TADs were called with hicFindTADs ( --minDepth 15000 --maxDepth 150000 --step 15000 --thresholdComparisons 0.05--delta 0.01 --correctForMultipleTesting fdr) from the HiCExplorer suite (60).

#### Differential transcript usage

Mouse RNA-Seq was quantified with Salmon (64) against our previously published BMDM transcriptome (52). To reduce complexity, representative gene models were selected as in Table S8 to represent the isoforms of *Loup* and *Spi1*. TPMs were imported with Tximport (65) and relative transcript usage calculated with DRIMSeq (66) by treating the Loup, Spi1_AFE1 and Spi2_AFE2, all of which begin at the upstream exon, as a single feature.

#### Ribo-Seq

Adapters were removed with Cutadapt (49). A tRNA/rRNA index was built with Bowtie2 (59) and mapped reads were discarded. The remaining reads were mapped with STAR in end-to-end mode to the Hg38 genome, guided by a custom annotation set composed of Gencode v41 merged with the published isoform atlas from primary macrophages (27). Multimapping reads were discarded.

### Western Bloting

Cell lysates were quantified by the Pierce^TM^ BCA Protein Assay Kit. Equal amounts of protein (15 ug) of each sample were denatured at 70°C for 10 min prior to loading on 12% SDS-PAGE. Samples were transferred to polyvinylidene difluoride (PVDF) membranes using the Trans-Blot Turbo Transfer System (Bio-Rad), blocked with TBST (1x Tris buffered saline with 0.1% Tween 20) supplemented with 5% (wt/vol) BSA for 1 hr and blotted with PU.1 (9G7) (1:1000, Cell Signaling #2258) at 4°C overnight. Horseradish peroxidase (HRP)-conjugated goat anti-rabbit (1:2000, Bio-Rad, #1706515) secondary antibody was used. Western Blots were developed using SuperSignal™ West Pico PLUS Chemiluminescent Substrate (Thermo Scientific cat# 34577). After imaging HRP was inactivated for 2 hours in 0.2% Sodium Azide in TBST supplemented with 5% (wt/vol) BSA. B-Actin (C4) monoclonal antibody (1:500, Santa Cruz Biotechnology cat #47778) was subsequently used as a loading control followed by HRP-conjugated goat anti-rabbit secondary antibody (1:2000).

### siRNA knockdown of SPI1

WT THP1 cells were transfected with 60 pmol of SPI1 targeting (ThermoFisher cat# HSS186060) or Cy3-conjugated non-targeting siRNA (ThermoFisher cat# AM4621) for 72 hours using Lipofectamine 3000 (ThermoFisher cat# L3000001) according to the manufacturer’s protocol.

### Nanostring multiplexed transcript analysis

RNA was isolated using the Direct-zol RNA Miniprep Plus Kit (Zymo cat# R2052) from one control and two *LOUP* knockdown THP1 cell lines at baseline and after 6 hours of LPS treatment. RNA was quantified on a Nanodrop. 100ng of RNA was used for each sample hybridization (Nanostring Master Kit cat# 100052) and detection with the Nanostring Human Immunology V2 nCounter GX Codeset (cat# 115000062) on a MAX/Flex nCounter according to the manufacturer’s protocol.

### ELISA and Multiplexed ELISA

WT THP1 cells transfected with SPI1 targeting siRNAs were seeded at equal densities and stimulated with LPS for 24 hours. Samples were diluted 1:10 and IL8 was measured using the DuoSet IL8 ELISA kit (R&D Systems cat# DY208) following the manufacturer’s protocol. THP1-NFkB-EGFP-CRISPRi-sgRNA *LOUP* knockdown and control cells were seeded at equal densities and treated with LPS for 24 hours. 75ul of supernatants were collected and analyzed undiluted by EVE Technologies using their Human Cytokine Panel A 48-Plex Discovery Assay (cat# HD48A).

### Nuc/Cyt fractionation and RT-qPCR

WT THP1 cells were fractionated according to the NE-PER kit protocol (ThermoFisher cat# 78833) with RNAse inhibitor (Superase-IN, ThermoFisher cat# AM2696) added to the cytosolic and nuclear lysis buffers. 3 volumes of Trizol (TRI Reagent, Sigma T9424) was added to the fractions and RNA was isolated using the DIrect-zol RNA Miniprep Plus Kit (Zymo cat# R2052). 16uL of RNA isolated from fractions was reverse transcribed (iScript cDNA synthesis kit, Bio-Rad cat# 1708840) followed by qPCR (iTaq SYBRgreen Supermix, Bio-Rad cat# 1725121) using the cycling conditions as follows: 50C for 2 min, 95C for 2 min followed by 40 cycles of 95C for 15s, 60C for 30s and 72C for 45s.

### RNA Isolation and RT-qPCR of Primary Mouse Monocytes

RNA was extracted from primary mouse monocytes following treatment with 200 ng/mL LPS for 2 or 6 hours or no treatment. RNA was isolated by phenol chloroform extraction and ethanol precipitation as follows: RNA sample mixed in 5:1 ratio of tri-reagent (Sigma Aldrich, T9424-200 mL) to chloroform (Ricca, RSOC0020-500C), sample spun max speed for 20 mins at 4C in gel-heavy tubes. The upper aqueous layer was removed to a new tube and mixed 1:1 with Isopropanol (Fisher Scientific, BP2618-4) and incubated for 1 min with glycoblue co-precipitant (Invitrogen, AM9515). Sample was spun at 5 min at max speed at 4C. Supernatant was discarded, and pellet was washed with cold 75% Ethanol and spun at max speed at 4C for 2 mins 2X, discarding supernatant between spins. Pellet was air dried for 1 min. Sample subjected to DNAseI treatment (New England Biolabs, M0303S) at 37C for 30 mins. After DNase treatment added 2X NT2 buffer and equal total volume of lower layer of Phenol/Chloroform/Isoamyl Alcohol (Fisher Scientific, BP17521-400), vortexed, spun room temp 5 min at max speed. Upper aqueous layer was transferred to a new tube and mixed with Sodium Acetate to a final concentration of 0.3 M, 2.5X 100% EtOH, and glycoblue co-precipitant prior to precipitation at -80C for 1-3 days. Sample spun at max speed for 30 mins, supernatant discarded. Pellet washed with 75% EtOH and spun 2 mins at 4C 2X, discarding supernatant in between. Pellet air dried RT 2 mins and resuspended in DEPC water.

RNA quantified using Qubit RNA HS kit (Invitrogen, Q32852). Equal amounts of RNA per replicate (250-560 ng) was reverse transcribed into cDNA using iScript Reverse Transcription Supermix reagent (BioRad, 1708890). qPCR was performed using primers against murine HPRT and murine *LOUP* and iTaq Universal Sybr Green Supermix(BioRad, L001751B) with the following cycling conditions 95C for 10 min, followed by 41 cycles of 95C for 15s, 60C for 30 s, 72C for 30s, followed by 95C for 10s and a melt curve from 55 to 95C at 0.5°C per second. Primer sequences for mouse found in supplemental table 1. Gene expression was normalized to HPRT.

**Tables 1 and 2-Description of top hits from NFkB and PMA screen.**

Each significant hit gene and its closest protein-coding neighbor are described. Data from the Genome CRISPR database was used to note whether the targeted lncRNAs or their neighbors have been previously indicated as hits in viability screens, if so they have been labeled “viability hit”. Four of the hit genes once annotated as lncRNAs have been updated to protein coding genes and therefore their neighboring genes were not assessed. Distances between targets and their neighbors were measured between promoters predicted by the ENCODE Registry of Candidate Cis-regulatory Elements on the UCSC genome browser.

**Supp** **Fig.1** **-Tissue expression of *LOUP*, *SPI1* and neighboring genes.**

**A. Gene models at *LOUP*/*SPI1* locus.** *SPI1*, *LOUP*, and *SLC39A13* gene models were determined by longread R2C2 and Mandalorian from primary macrophages. *MYBPC3* is the Gencode v44 gene model.

**B-E. GTEx tissue expression of genes.** B. *MYBPC3* is primarily expressed in heart tissue. C. *SLC39A13* is highly expressed in multiple tissue types. D.,E. *LOUP* and *SPI* are expressed primarily in whole blood.

**Supp Fig.2 - Sequence and structural conservation of LOUP from human and mouse**

A. **Sequence alignment of transcripts.** Emboss Needle pairwise alignment of human and mouse *LOUP* transcripts. B,C. Vienna Fold structure predictions of human and mouse *LOUP*.

**Supp Fig.3 - Differential transcript usage after LPS**

**Mouse *Loup* decreases following LPS**. BMDMs stimulated with LPS for 6 hours show relative decrease in expression of *Loup* while usage of *Spi1* alternate first exon isoforms is unchanged. Significance determined by DRIMSeq adj. p-values.

## Notes

### Competing Interest Statement

Susan Carpenter is a paid consultant for NextRNA Therapeutics

