## Supplementary figures and images for "CRISPRi screens identify the lncRNA, *LOUP,* as a multifunctional locus regulating macrophage differentiation epigenetically and inflammatory signaling through a short, encoded peptide"

### Supp_Fig.1

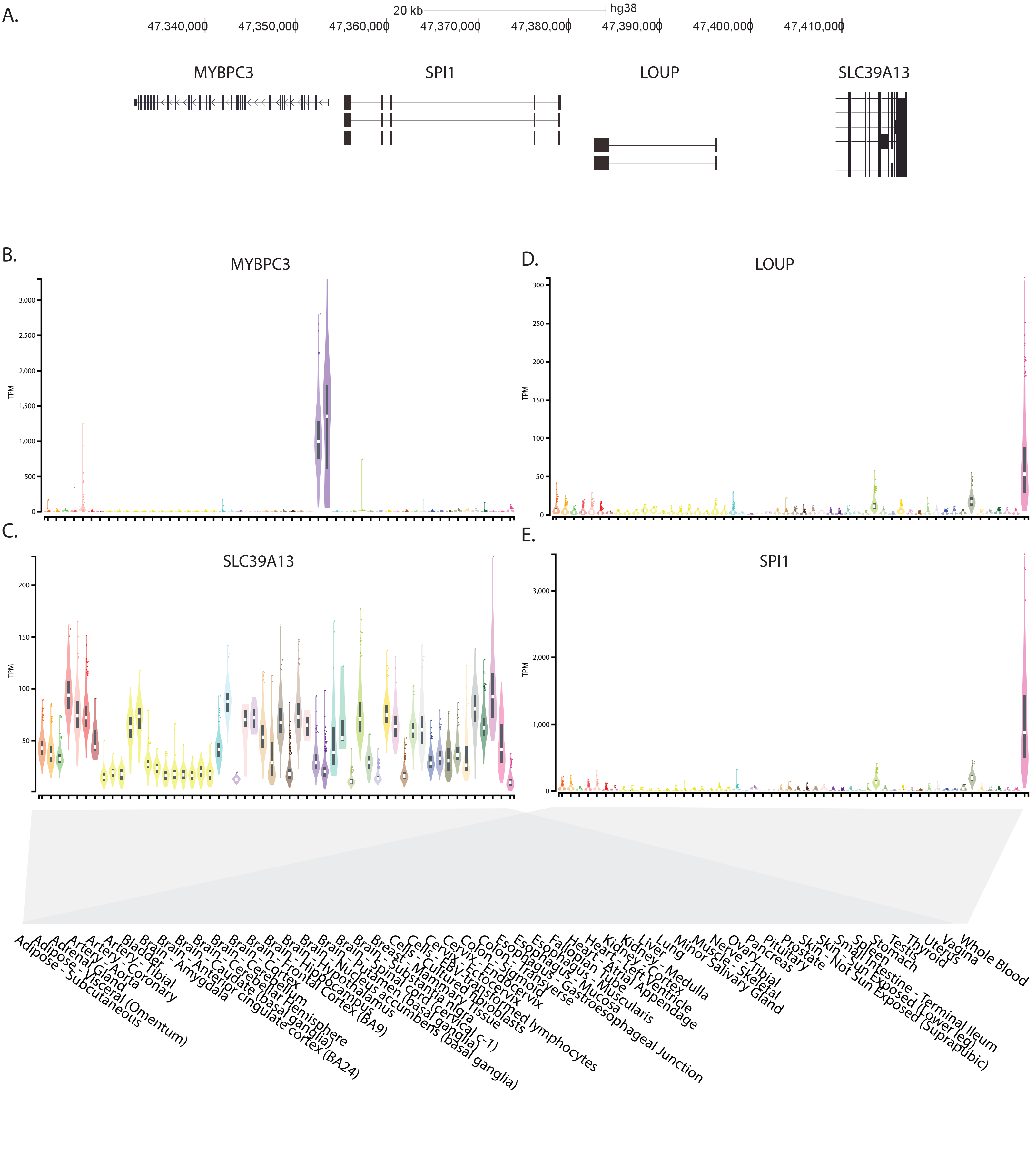

### Supp_Fig.2

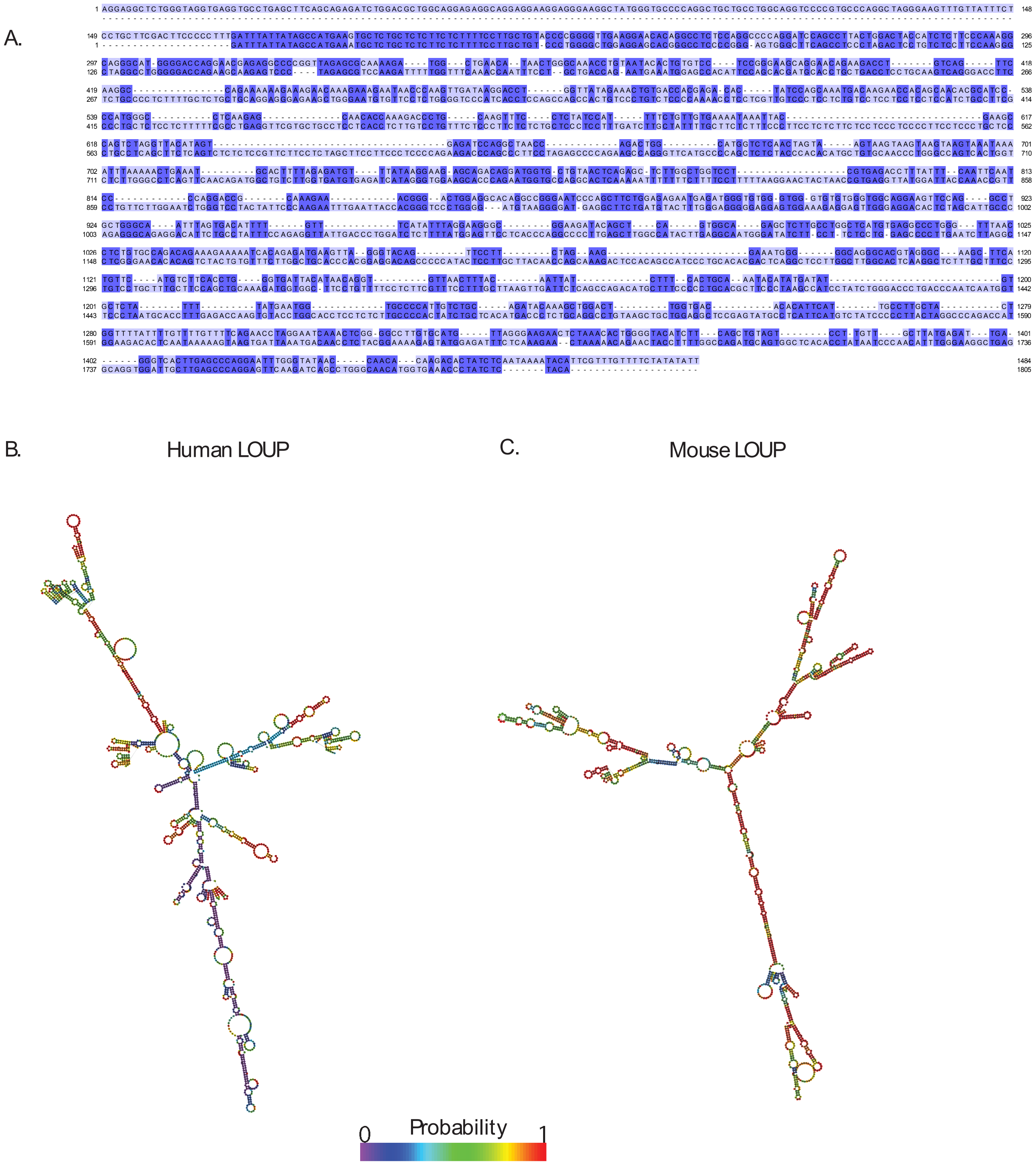

### Supp_Fig.3

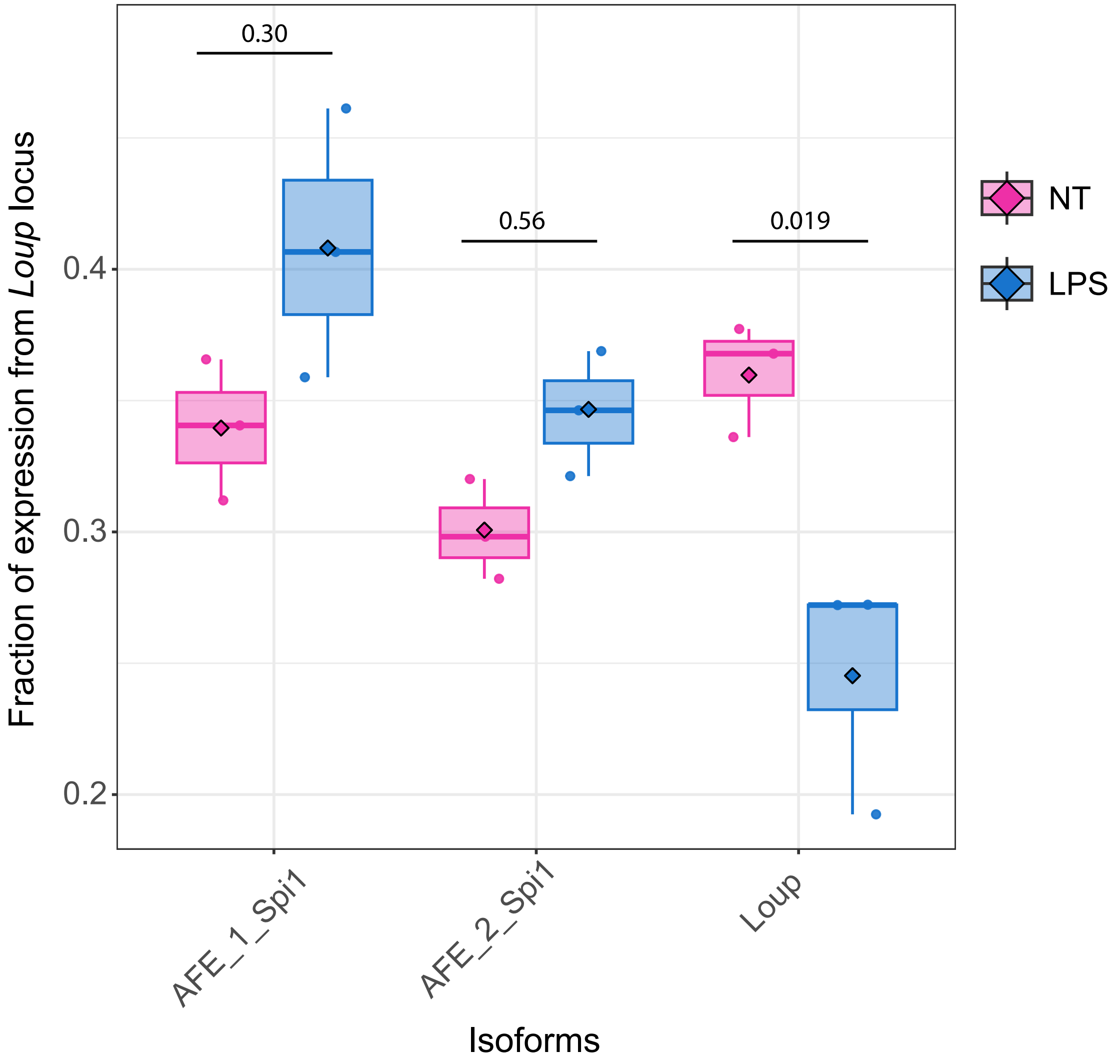
